# Machine learning guided batched design of a bacterial Ribosome Binding Site

**DOI:** 10.1101/2022.01.05.475140

**Authors:** Mengyan Zhang, Maciej Bartosz Holowko, Huw Hayman Zumpe, Cheng Soon Ong

**Affiliations:** Machine Learning and Artificial Intelligence Future Science Platform, CSIRO; Department of Computer Science, Australian National University; Data61, CSIRO; Synthetic Biology Future Science Platform, CSIRO; Land and Water, CSIRO

**Keywords:** machine learning, optimisation, genetic part design, ribosome binding site

## Abstract

Optimisation of gene expression levels is an essential part of the organism design process. Fine control of this process can be achieved through engineering transcription and translation control elements, including the ribosome binding site (RBS). Unfortunately, design of specific genetic parts can still be challenging due to lack of reliable design methods. To address this problem, we have created a machine learning guided Design-Build-Test-Learn (DBTL) cycle for the experimental design of bacterial RBSs to show how small genetic parts can be reliably designed using relatively small, high-quality data sets. We used Gaussian Process Regression for the Learn phase of cycle and the Upper Confidence Bound multi-armed bandit algorithm for the Design of genetic variants to be tested in vivo. We have integrated these machine learning algorithms with laboratory automation and high-throughput processes for reliable data generation. Notably, by Testing a total of 450 RBS variants in four DBTL cycles, we experimentally validated RBSs with high translation initiation rates equalling or exceeding our benchmark RBS by up to 34%. Overall, our results show that machine learning is a powerful tool for designing RBSs, and they pave the way towards more complicated genetic devices.

## 1 Introduction

One of the main tenets of synthetic biology is the design, evaluation and standardisation of genetic parts [1, 2, 3]. A central challenge is part design, understood as modifying the sequence of a genetic part for it to meet specific requirements. Genetic parts are the units which are ultimately combined into more complex genetic circuits that produce desired functions in the target organisms. This is usually done in terms of the Design-Build-Test-Learn (DBTL) cycle, where the given genetic part or organism is continually improved through an iterative process. This cycle involves designing (D) new DNA sequences to achieve a desired property in computer-aided design software, then physically building (B) new constructed variants and testing (T) using an analytical instrument in a laboratory. Computer modelling can be used to learn (L) and predict characteristics of a genetic part [4, 5]. Most of these computer models are based on either the thermodynamic properties of the involved molecules (DNA, RNA, proteins, etc.) or empirically-obtained values describing a relevant design property, like translation initiation rate (TIR) in the case of ribosome binding sites (RBSs) [6, 7, 8]. However, de novo design of small genetic elements is still challenging due to unknown relationships between their sequence and performance.

In this paper, we propose a machine learning guided Design-Build-Test-Learn cycle for the experimental design of bacterial RBSs. This consists of two distinct types of machine learning methods, one in the Learn phase and a second in the Design phase. We show how small genetic parts can be reliably designed using even relatively small, but high-quality data sets. This work focuses on RBS part design and TIR prediction, rather than looking at its wider genetic context and impact on the general performance of the cell. As the RBS is one of the key genetic elements controlling protein expression in bacteria, it is a suitable target for establishing a workflow that could be potentially translated to more complicated systems.

In the Design phase of the DBTL cycle, designers often have to fine tune characteristics of parts to give the resulting strains their desired properties. The ability to predict a characteristic of a genetic part (for example using a machine learning method) only provides inputs to the problem. The designer still needs to choose from the large number of possible variants to Build. For instance, if the goal is to increase the yield of a protein, the designer might increase the TIR of the RBS responsible for translation of that protein. Hence the goal for the Design phase of TIR is to select a small set of RBS sequences from the design space (i.e. all possible DNA sequences) based on the predictions from Learn phase. However since the computational predictions are not perfectly accurate, there is uncertainty about which potential RBS sequence has the desired property (for example highest protein yield). The designer could use the mean predicted TIR values and choose the best ones to exploit the knowledge modelled by the computational predictions. But the designer may also want to explore regions of DNA design space where the predictions are highly uncertain (e.g. as measured by the predicted TIR standard deviation) and could yield high payoff. Hence a major challenge of the Design phase is to balance the exploitation and exploration, which is addressed by one class of machine learning approaches called multi-armed bandits [9]. As we will see in this paper, multi-armed bandits are well suited to solving the challenge of recommending a small set of RBS sequences in the Design phase of the DBTL cycle.

One way to generate predicted TIR values is to use the existing RBS calculators. Reeve et al. [8] surveys three main RBS calculators, all predicting the TIR based on the thermodynamic properties of the RBS and the ribosome [10, 11, 12]. Reported predictions from all of these models range from relatively good (*R*^2^ > 0.8) to low (*R*^2^ < 0.2) depending on the data set [13]. This may be due to number of reasons: i) they rely on calculations of free energies which can be difficult to estimate with high precision, ii) in general, one of the best ways to improve the models’ accuracy is by increasing the number of phenomena taken into account, which can lead paradoxically to decreased model accuracy due to accumulation of errors [14], and iii) by using deterministic coefficients to calculate energies, one disregards the often stochastic nature of processes in cells which can potentially increase prediction error [15]. There is also evidence that binding energy calculations may be poor predictors of RBS strength [16, 17]. This is reinforced by studies suggesting that RNA secondary structure is potentially a more important feature in TIR determination [18, 14]. This suggests that multiple interactions determine the mRNA-ribosome binding, and predicting TIR from the genomic sequence is still challenging.

Recent work has explored the use of machine learning predictors to learn from historical data and generate predictions for use in synthetic biology, vastly improving the DBTL cycle’s performance [19, 20, 21]. This leverages the exponential increase in experimental data from synthetic biology [22] to improve predictors in the Learn phase of the DBTL cycle. For example, Jervis et al. [23] used support vector machines and neural networks to optimise production of monoterpenoids in *Escherichia coli*. Similarly, Costello et al. [24] have used a number of machine learning approaches to analyse time-series multi-omics data to predict metabolic pathway behaviour. Deep learning techniques have also been successfully used to analyse large synthetic biology data sets [25, 26, 27]. Recall that the multi-armed bandit approach balances the exploration-exploitation trade off by using predictive uncertainty. The machine learning predictors mentioned, as well as existing RBS calculators, do not yet provide uncertainty levels to guide exploration. Furthermore large data sets may not be available for a genetic part of interest, for instance the RBS. Hence there is a need to develop methods for training predictors on smaller data sets and generate predictions for both mean values and uncertainties in the Learn phase of DBTL. The predictions of characteristics of interest for each potential variant can then be used in the Design phase.

Our overall experimental goal is to maximise the TIR by building and testing batches of RBS sequences with only a small number of DBTL cycle iterations. We demonstrate how machine learning (ML) algorithms can be incorporated into the DBTL cycle to predict (Learn) and recommend (Design) variants of a bacterial RBS with the goal of optimising associated protein expression level in the specific genetic context (i.e. with specified bases upstream and downstream of the investigated RBS sequences). Our proposed machine learning guided DBTL cycle is summarized in Figure 1. Two types of ML algorithms are applied in the DBTL cycle. For the Learn phase, the goal is to train a predictor that will predict the protein expression level by learning from the logged data. We used a Bayesian, non-parametric regression algorithm, called Gaussian Process Regression (GPR) [28] in our pipeline. In addition to providing a predicted mean TIR, GPR also provides uncertainty estimates via its predicted standard deviation. GPR has been shown to perform well with small amounts of trainig data in biological prediction tasks [28, 29]. For the Design phase, the goal is to find a policy to recommend RBS sequences to query in batches so that we can identify the optimized RBS sequences within a given budget. We use a version of the Upper Confidence Bound multi-armed Bandit algorithms [29] to learn the policy. The policy uses the outputs of the GPR predictions to recommend useful batches for the Build and Test phases. We demonstrate that laboratory automation methods in the Build and Test phases result in high quality TIR data, that is well suited for training machine learning methods in the subsequent Learn phase. The two types of algorithms cooperate with each other and provide powerful tools for DBTL cycle in genetic parts design [30, 29]. Using our proposed machine learning guided DBTL cycle (Figure 1), we were able to increase expression of our target protein by up to 35%, as compared to the very strong benchmark RBS.

**Figure 1:**
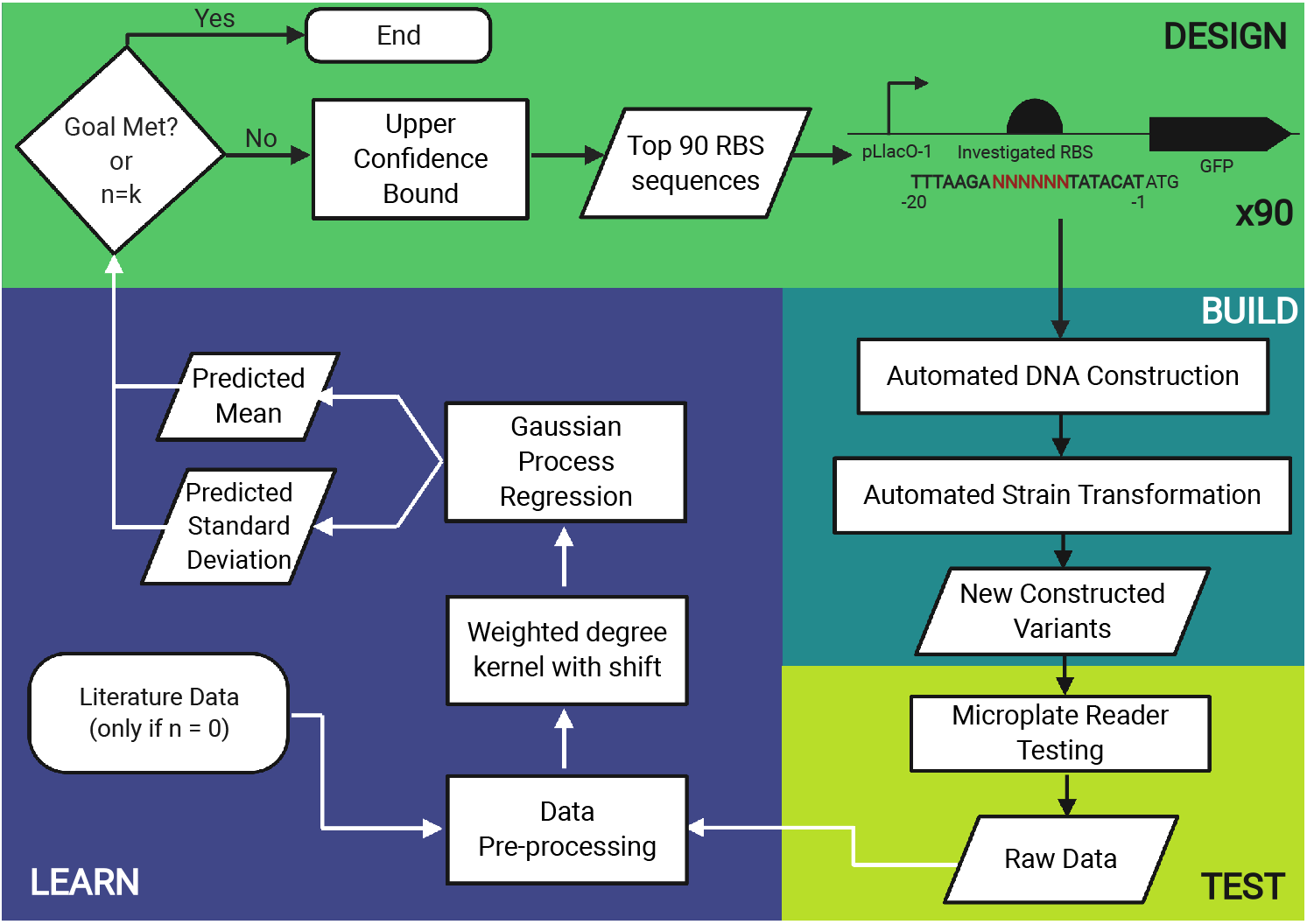
Flowchart of the machine learning-based experimental design. The RBS design is recommended by the Upper Confidence Bound multi-armed Bandit algorithm. After generating the recommendations, the RBSs are built and tested using automated laboratory methods allowing for rapid construction and testing at scale. Finally, the obtained results are fed back to the Gaussian Process Regression prediction algorithm in the Learn phase. n is the current design round and k is the maximum number of rounds allowed by time and/or money. In regards to the “Goal Met?” condition, the goal in our case was to find sequences with TIR significantly higher than benchmark, but the goal can be generalised to fit the requirements of the user.

## 2 Results

We present our RBS-optimising DBTL workflow that uses machine learning in Section 2.1. Machine learning is used in two different ways: i) we show the efficacy of the ML recommendations in the Design stage (Section 2.2), and ii) we demonstrate that the ML predictions are accurate in the Learn stage (Section 2.3). We present our new RBS sequence library in Section 2.4 and describe some interesting characteristics of the discovered sequences, as well as show the effectiveness of the automated laboratory workflow.

### 2.1 The experimental workflow

Our DBTL workflow, which uses machine learning to optimise protein expression, is shown in Figure 1. Build and Test are driven chiefly by choices made by human researchers and the use of automated methods. Machine learning algorithms are applied in Learn and Design. In the Learn phase, we use the Gaussian Process regression algorithm to predict the TIR of RBS sequences comprising the experimental space. In the Design phase, we use the Bandit algorithm to recommend new RBS sequences based on the predictions from Learn.

The exact position of the RBS in the sequence upstream of the protein coding sequence (CDS) can be hard to pinpoint, however most previous studies put it in the 20 bp directly preceding the coding sequence. In our investigation, we are using an RBS which is known to have a very high TIR when expressing GFP and is present in the pBb series of plasmids [31]. This template RBS sequence is 20 base pairs (bp) long with the sequence TT-TAAGA**AGGAGA**TATACAT, where the highlighted nucleotides constitute the core of the RBS. Since this is the sequence against which new RBS sequences will be benchmarked, we will refer to this sequence as the *benchmark sequence* hereafter. Additionally, we have experimentally confirmed that modifying the core sequence is statistically more impactful on TIR than changes made outside of it (see Figure S2). This hypothesis has been built based on reported biases towards certain bases present in the core of the RBS but absent outside of it. For example, according to [32] there is a strong bias towards A and G bases in the core region of the RBS. Similarly, outside of the 6 bases of the core in the wider 20 bp context of the RBS there is no significant bias towards any particular base which suggest that these bases do not contribute to the overall TIR of a given RBS. Focusing on the 6 to 8 bp core sequence is a common RBS design approach [33]. Hence, in our design, we focus on modification of the core at nucleotide positions −8 to −13 (relative to the start codon of the GFP; this is where the consensus Shine-Dalgarno AGGAGG sequence is usually found in wild type *E. coli*) of the RBS and we keep other positions the same as the benchmark sequence, i.e. TTTAAGA + NNNNNN + TATACAT, where N can be any nucleotide (A, C, G, T). The total experimental (variant) space is then 4^6^ = 4096.

In our genetic design, the investigated RBS controls expression of green fluorescent protein (GFP). By controlling expression of a fluorescent protein with the RBS we can quickly assess the perceived relative TIR by measuring fluorescence of cells harbouring the expression vector over time. Finally, the mRNA is transcribed from an IPTG-inducible promoter pLlacO-1. Inducible expression allows one to synchronise the start of the GFP expression in all the cultures with addition of IPTG. Since standardisation and comparative studies should be done in as similar genetic context as possible, the design of this device has been deliberately kept simple to make such studies easier [34].

### 2.2 DESIGN: Performance of the recommendation algorithm

The Bandit recommendations were made using the batch Upper Confidence Bound multi-armed Bandit algorithm [29]. In short, this algorithm is a stochastic method of probing the experimental space, and is sometimes referred to as Bayesian optimization. It aims at maximising the reward (output) from testing a limited number of instances from a large pool which cannot be tested exhaustively due to limited resources (time, computational power, money). It balances the exploration-exploitation paradigm, where exploration focuses on testing data points which maximise information gain and exploitation focuses on recommending RBSs with high predicted TIR.

Figure 2A shows the results for all the RBS groups tested experimentally. In each experimental round, in addition to the new RBS designs, we measure the TIR of the benchmark RBS as the internal standard. We then obtain the normalised TIR (called *TIR ratio*) by taking the ratio between the raw TIR of a new design and the average TIR of benchmark sequences in each round (which are run in triplicate in each round). Figure S3 shows these results in terms of raw TIRs.

**Figure 2:**
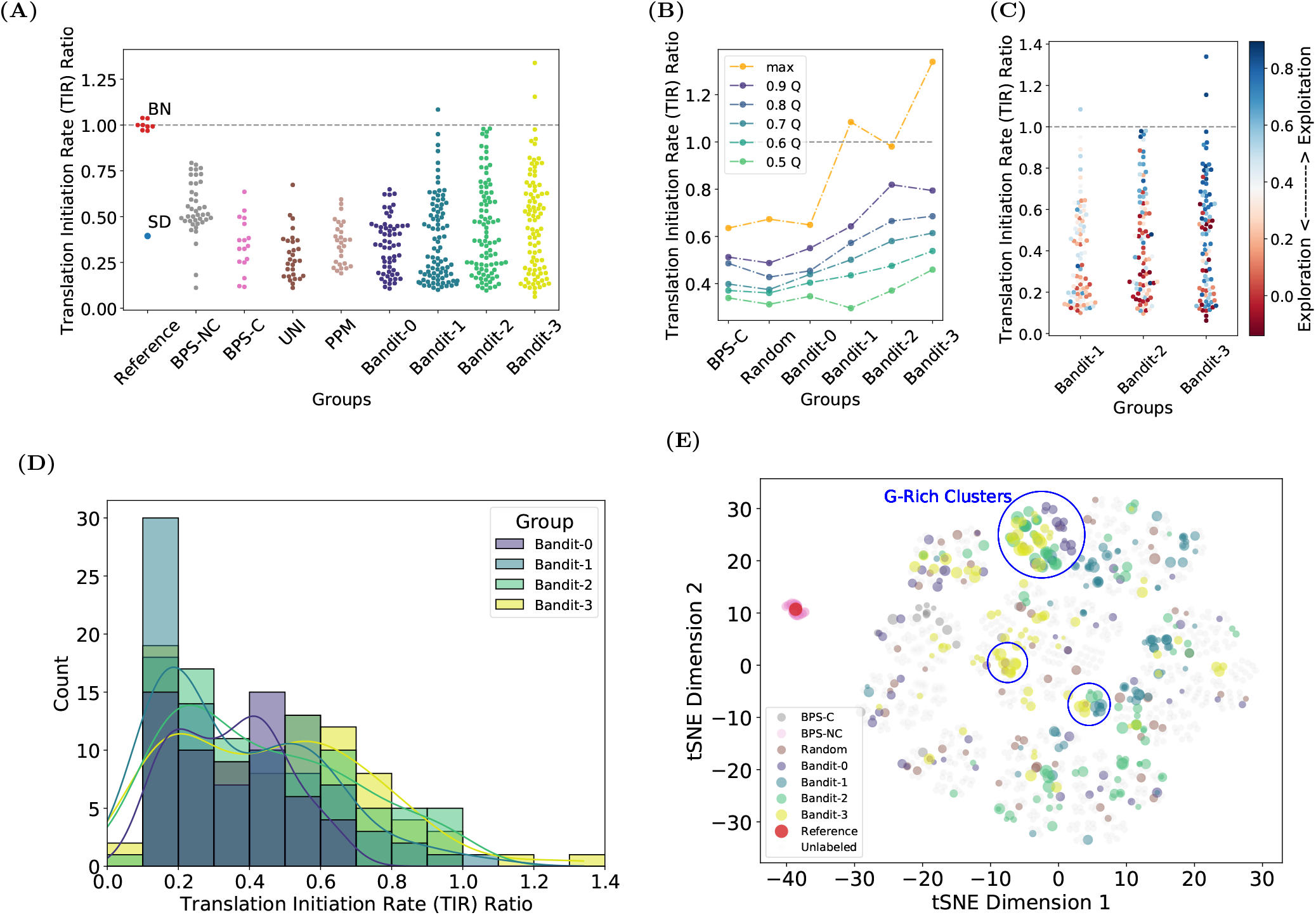
TIRs of RBS groups examined in this study. **A)** Swarm plot showing the obtained TIRs divided into RBS groups. BPS-NC: base-by-base changes in the non-core region. BPS-C: base-by-base changes in the core region. UNI: Randomly generated sequences with uniform distribution. PPM: Randomly generated sequences with distribution following the Position Probability Matrix for all natural RBS in *E. coli*. Bandit-0/1/2/3 - Bandit algorithm generated results for Round 0, 1, 2 and 3 respectively. SD - Shine-Dalgarno sequence. Dashed line is set to 1 and represents the averaged benchmark sequence TIR for that group. BN - benchmark sequences for all plates. (not all are exactly 1 due to them being shown as separate samples rather than per round averages.) **B)** Line plot showing TIR obtained in a given quantile (Q) of results divided into groups as in A). UNI and PPM are merged into Random group and BPS-NC is not shown due to changes being made outside the core in that group. **C)** Exploitation v.s. Exploration for Bandit 1-3. Blue-hued points represent exploitation, those hued red represent exploration. **D)** Histogram with kernel density estimations (KDE) showing distributions of TIRs for Bandit groups. **E)** t-SNE plot showing the relative distances between sequences in our design spaces as calculated by our kernel function (weighted degree kernel with shift). The area of the circle corresponds to the experimentally-obtained TIR value. The TIR results in all subplots are shown normalised to the respective benchmark sequence sample which acts as internal standard; the TIR of a given RBS is divided by TIR of the benchmark RBS run in the same plate.

To generate the data set from which the algorithm would learn, we decided to characterise a total of 450 RBS variants, little over 10% of the whole experimental space. To fit our automated workflow, we divided the 450 variants into batches of 90, split into 4 design rounds.

In the zeroth Round we tested two batches of designs, giving a total of 180 variants split as below:

- BPS-NC and BPS-C group: 60 RBS sequences which are subsequent single nucleotide variations of all 20 nucleotides of the original, benchmark sequence. This batch is designed to show us influence of such single nucleotide changes on the overall performance of the RBS and the potential impact of changes made beyond the core part (see Supplementary Figure S2).
- UNI group: 30 RBS sequences that were uniformly randomised, i.e. equal probability of choosing any nucleotide for each position. This group shows the performance of RBSs generated randomly.
- PPM group: 30 RBS sequences randomised based on the position probability matrix (PPM) generated from all the naturally-occurring RBS sequences in the *E. coli* genome [35]. This group shows the performance of RBSs generated randomly, but following the natural nucleotide distribution.
- Bandit-0: 60 RBS sequences recommended by our implementation of the recommendation algorithm based on a data set obtained from literature [36], which contains 113 non-repeated records for 56 unique RBS sequences with their respective TIRs. This data set has been used due to the perceived similarity of its goal to that of this work - prediction of TIR based on phenotypic output.

In the subsequent 3 rounds, with one batch each, all 90 designs were generated using our machine learning algorithm based on the data obtained from the previous rounds (these groups are called Bandit 1 to 3 respectively).

All Round 0 groups (BPS-NC, BPS-C, UNI, PPM, Bandit-0) performed worse than our benchmark sequence in terms of TIR. The best-performing group was the BPS-NC, which is explained by the relatively small impact on the TIR of changes made outside the RBS core. The Bandit-0 group’s performance is similar to randomly generated designs, despite being machine learning-driven, due to being trained on approximate data. Starting from Round 1, where the prediction and recommendation algorithms were fed data from Round 0, the results improved significantly, with a number of sequences performing better than the consensus Shine-Dalgarno sequence and in one case, better than the benchmark (by 8%). In Round 2 we observed further improvement by obtaining more sequences with TIRs similar to our benchmark sequence. Finally, in Round 3 the algorithm identified two sequences that were 34% and 15% stronger than the benchmark sequence.

In summary, out of 450 tested sequences, around 40% (BPS-NC, BPS-C, UNI, PPM) were created through some kind of sequence randomisation. Out of these randomised sequences, only a few got close to the benchmark sequence’s TIR and were still 20% weaker than it is (Figure 3A). In fact, these 80% TIR ratio sequences were created by randomising the sequence outside of the core RBS region (BPS-NC), which was statistically shown not to be significantly impactful on the TIR (see Supplementary Figure S1 and, for example, work by Jeschek et al. [33]). More representative would be sequences from other random groups (BPS-C, UNI, PPM), from which the best sequence achieved only about 65% of the benchmark TIR. These results show that generating a strong RBS sequence by random mutation is a non-trivial task, when the tested data set is relatively small. Contrasted with this, non-randomized, our Bandit-driven design batch gave much better results, with RBSs getting close to benchmark performance and even exceeding it.

**Figure 3:**
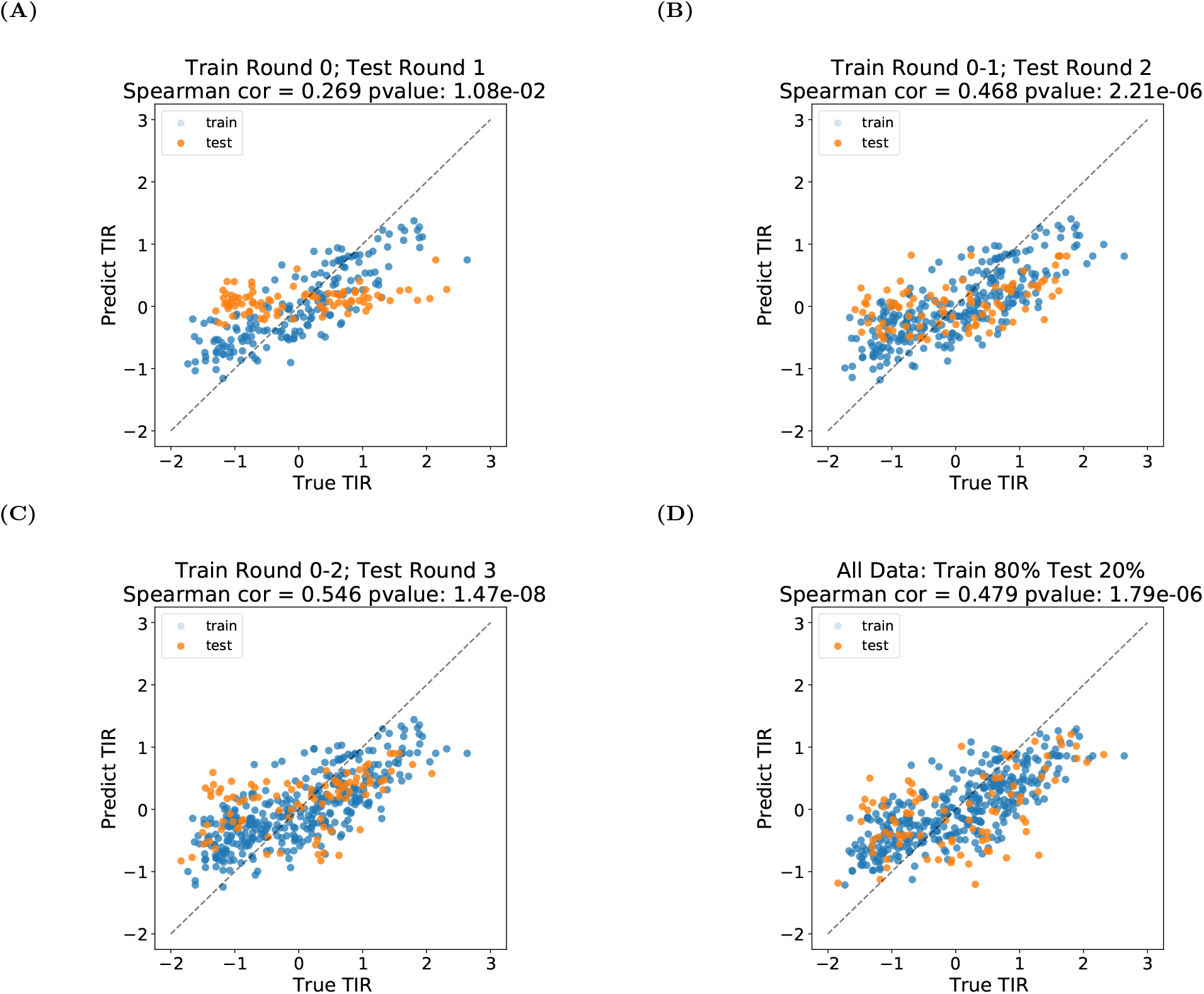
Performance of the prediction algorithm (no kernel normalisation). The scatter plots A-D show the performance of our prediction algorithm calculated after each round. Note that the TIR values are normalised according to the standardisation described in section 4.2.1, which is different from the TIR ratio reported in Figure 2. X-axis and y-axis are the true, tested TIR and the predicted TIR respectively. In (A)(B)(C), we show at round *t* = 1, 2, 3 respectively, we train our predictor based on the previously obtained data (round 0 to *t* − 1) and show the predictions on both the training data (orange) and recommendations suggested by the Design phase (i.e. test data, blue). In D), we train our predictor on randomly chosen 80% of all available data, and test on the remaining 20% data. The Spearman correlation coefficient (with corresponding p-value; calculated using test data only) are provided in each plot’s title on test data only. The p-value here is for the null hypothesis stating that two sets of data are uncorrelated.

Figure 2B shows the same results but divided into quantiles where the specific point for a given group shows the highest TIR for that quantile. The gradual increase for all quantiles can be observed for all Bandit groups, suggesting that the algorithms have a better understanding of the experimental space given more data. The decreased result in the 0.9th quantile compared to the maximum value for the Bandit 3 group can be attributed to the increased emphasis on exploitation that has been set for that round compared to others. We see this effect in Figure 2C (with details shown in Supplementary A.6 and Figure S1), where we coloured the data points for Bandit 1-3 groups according to their relative exploration - exploitation affinity. Those with a high predicted mean are coloured blue and represent exploitation, those coloured red are with high predicted uncertainty and represent exploration. The implication of the fact that an RBS sequence chosen at random will have low TIR (as shown by UNI and PPM), is that most of the exploration will result in low TIRs. Our results confirm that RBSs with high TIRs tend to come from exploitation of the design space, whereas the exploration points give relatively low TIRs. Note that the exploration is necessary to expand our knowledge to the unknown parts of the design space and in effect allow us to exploit it better.

Figure 2D shows the TIRs of RBSs tested in the Bandit groups divided into bins with width equal to a TIR ratio of 0.1. KDE plots have been overlaid to depict the calculated density for each group. The increase in prevalence of later Bandit groups in the higher bins is evident, especially for Bandit 2 and 3, constituting the bulk of results in the > 0.8 TIR ratio bins. Notably, the distributions calculated for all the groups are bimodal - we discuss the possible reasons for that later in the text.

In Figure 2E we show a t-distributed stochastic neighbour embedding (t-SNE) [37] plot depicting the experimental space. Each RBS is located on the plot according to its distance from other RBSs as calculated by our weighted degree kernel with shift (see Section 4.2.2). The RBSs recommended by Bandit groups have covered the majority of the design space. Additionally, a number of clusters were especially targeted by our recommendation algorithm. For example, the circled clusters labelled as “G-Rich Clusters” have been actively recommended by the algorithm. More specifically, sequences with 4 or more guanines in any position constituted 10% of the randomly selected sequences and 5, 9, 16 and finally 25% in each of the 4 Bandit guided batches respectively.

### 2.3 LEARN: Prediction of RBS performance

To provide the predicted TIR with the confidence interval needed for the UCB algorithm in the Design phase, we have selected a regression algorithm, Gaussian Process Regression (GPR), which can predict not only the means, but also uncertainty estimations for our data points. Figure 3 shows how our implementation of the Gaussian Process algorithm performed in terms of predictions in each round. As expected, the predictions in Round 0 were poor due to the use of approximate data. The predictions improved for the subsequent rounds, the Spearman correlation coefficient rose from 0.269 for Round 0 to 0.546 for Round 3.

The regression task in the DBTL cycle is more challenging than the large-scale data-based regression tasks. In the early iterations, we have a limited number of data points and relatively high variability due to the measurement noise. However, since the predictions are only used for recommendations in the Design phase, instead of precise predictions of the mean for each RBS, we only need to provide a valid ranking for both the predicted mean TIR and uncertainties, and thus the ranking of the UCB scores. This would ensure that for each round we would be testing the required designs, even if we are not exactly accurate in terms of their numerical characteristics. Because of that, we provide the evaluation of the Learn phase in terms of a ranking-based Spearman correlation coefficient, which has been shown to be a more suitable evaluation metric when the prediction is used for recommendation tasks than coefficient of determination (*R*^2^) or Pearson correlation coefficient [38, 39].

Additionally, the performance of the Learn phase is also influenced by the exploration-exploitation balance in the Design phase. In each round, we select some data points for exploration of the areas in which we have few or none tested data points. For those exploration points, since the predictor never learns their label distributions and patterns, there is no chance for the predictor to provide accurate predictions of TIR values, thus hurting the prediction performance in the short term. However, this is useful information for future predictions as it allows us to understand the whole underlying space instead of focusing on local sub-optimal data points. In other words, we are intentionally sacrificing the accuracy of our predictions in each round to improve them in the future rounds. The effect of exploration of the space is the ability to find high TIR RBSs even with relatively low prediction performance.

### 2.4 BUILD & TEST: Characteristics of the tested sequences

We present some important characteristics of the tested RBSs in Table 1. Figure 4A shows the sequence logo calculated for the Top 30 sequences (Figure S5 shows the logo generated for all tested sequences). It is generally understood that guanine-rich sequences promote strong transcription. This expected bias towards guanine is clearly visible for all positions in our Top 30 RBSs. This result combined with the Bandit algorithm’s bias towards the G-rich cluster shown in Figure 2D reinforces the notion that our algorithm successfully identified G-rich sequences as the ones with high TIR probability.

**Table 1:** Characteristics of the library. Left table presents some of the characteristics of our library. Right table presents 10 RBS sequences with their corresponding TIR ratios; the first 9 are the strongest sequences including the benchmark sequence (AGGAGA) and the last is the Shine-Dalgarno sequence (AGGAGG). CV is coefficient of variation (standard deviation (STD) of a sample divided by its mean; see details in Figures S7 and S8). Efficiency of the Bandit design is calculated by dividing the highest TIR found using machine learning by the highest TIR found using random sequence generation.

| Characteristics of the library | Statistics |
| --- | --- |
| Total experimental space | 4096 |
| Planned constructs | 450 |
| Successfully constructed | 445 |
| Sequences with CV<40% | 79% |
| Sequences with CV<20% | 27% |
| Efficiency of Bandit design (compared with random) | 2 |
| Raw TIR range | [4.93, 105.38] |
| TIR ratio range | [0.06, 1.34] |

Table 1: **Characteristics of the library.**
| Top RBS Core | TIR Ratio |
| --- | --- |
| GGGGGC | 1.34 |
| GGGGGT | 1.15 |
| GGCTAT | 1.08 |
| **AGGAGA** | 1 |
| GGCGTT | 0.98 |
| GGGGGG | 0.98 |
| GGCGAC | 0.98 |
| CAGGAG | 0.96 |
| GGCGAG | 0.95 |
| **AGGAGG** | 0.39 |

**Figure 4:**
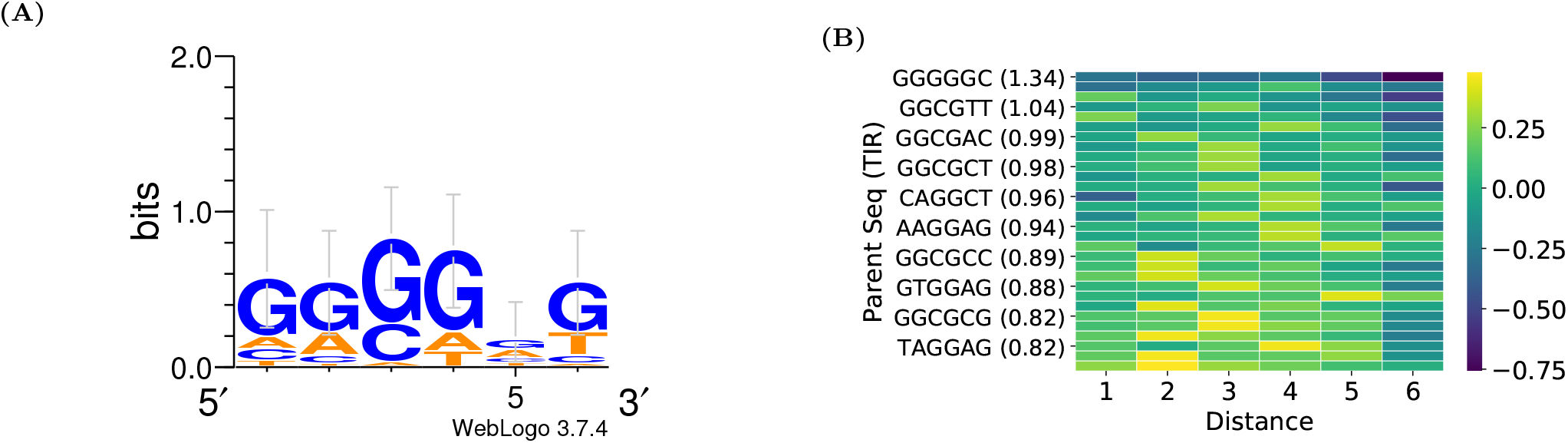
Characteristics of strong RBSs. A) Sequence logo calculated for the Top 30 tested sequences. B) Heatmap showing the edit (Hamming) distance required for positive change in TIR for RBSs with high and medium TIRs. The temperature scale shows the difference between a given RBS on the y-axis and the RBS with the strongest TIR at the given distance. Every second RBS is labelled for increased legibility.

Another interesting characteristic uncovered by our research is the perceived editing distance between two sequences required for improvement in the TIR when the given RBS’ TIR is already high. We define the edit distance as Hamming distance, that is, how many positions have to be changed to get from one sequence to the other (Hamming distance of 0 means that the sequences are identical and 6 means that they are two completely different sequences). Figure 4B shows the edit distance is required for positive change in TIR for an RBS with TIR > 0.75. For RBSs with high TIRs (> 1), the minimum distance that is required to increase the TIR is 2, with edit distances between 2 and 5 giving similar results. For RBSs with medium TIRs (< 1), a distance of 1 is enough to produce a meaningful increase in TIR.

This means that as the TIRs of examined RBSs increase, exploring sequences which are more dissimilar to the current candidates tends to give more meaningful improvement. As long as this does not impact targeted methods like machine learning-guided design, it does imply that the low rate of natural mutations will be very slow to explore more dissimilar sequences on such a short distance [40], which indicates that methods like Adaptive Laboratory Evolution may not be able to find very strong RBSs with a limited budget. In other words, because the examined sequence is relatively short (6 bp in a wider 20 bp context) the time required to accumulate 2 or more changes in the RBS region required for meaningful increase in TIR might be prohibitively long. In such cases, a directed process, like the one described here, should be strongly encouraged. This observation is in line with approaches seen in other disciplines, e.g. protein engineering, where more directed changes yield better results than wide random changes [41].

Finally, while our strong sequences showed some affinity for the anti-sense sequence of the ribosome known to bind to RBS, they did not show any obvious secondary structures that could explain their TIRs (see Figure S6). This result combined with the unexpectedly bimodal nature of KDEs in Figure 2 reinforces the notion, based on the previously reported literature [16, 14], that there may be a number of different mechanisms governing the probability of effective RBS-ribosome binding.

## 3 Discussion

In this work, we show how a machine learning guided approach, and high-throughput, automated laboratory methods can be jointly applied to efficiently optimise a small genetic part, in this case maximising the TIR of a bacterial RBS. In the Learn phase, we used Gaussian process regression to predict the TIR mean and standard deviation. To represent RBSs and capture the similarities between them, we choose to use the Weighted Degree Kernel with Shift method, which fits well with Gaussian processes. In the Design phase we used an Upper Confidence Bound multi-armed Bandit algorithm to recommend sequences to be tested in batches. In the Build and Test phase, we performed our experiments using laboratory automation to increase their speed, reliability and reproducibility. Using our proposed workflow and testing 450 RBS variants in 4 DBTL cycles, we designed and experimentally-validated RBSs with high translation initiation rates equalling or exceeding the currently known strong RBSs in this genetic context by up to 34%. Furthermore, we have generated an extensive library of diverse RBSs that can be used as a basis for future studies. In the rest of the section, we first revisit our overall experimental goal and challenges encountered in each phase of the DBTL cycle (Figure 1). We then link our proposed methods with related work and discuss the potential generalisation of our framework. We further discuss our design choices and open questions in this line of research. We refer [42] as more detailed discussion.

Our goal is to show the power of the machine learning guided DBTL cycle on RBS optimisation. Our approach has shown that this combination of the two machine learning algorithms is able to correctly detect and exploit rules of biological design that otherwise require substantial time and experiments to uncover. We focus on the part-centric optimisation [27], which is an important task in synthetic biology. Understanding how to design the individual parts also allows us to extend the framework to better strain design [23], where the part is optimised with a wider goal of strain optimisation.

The machine learning guided DBTL framework has good potential to be generalised to multi-gene pathway design as proposed by HamediRad et al. [43] and recently reviewed by Lawson et al. [21]. For example, the optimisation goal can be adjusted to address combinatorial optimisation for multi-gene and RBS scenarios; since this would be a large-scale data task, the current Gaussian Process regression prediction model could be updated to a deep Gaussian Process regression approach and the current Bandit algorithm could be optimised towards querying large design space to reduce computational complexity [44].

In the Design phase, we focus on maximising the TIR of RBSs, by gradually moving our emphasis from exploration to exploitation as we progress through the design rounds. While maximisation could be the appropriate payoff function for optimising RBSs or other small parts, other payoffs may be better in more complex cases. For example when considering multi-gene metabolic engineering, maximising expression of individual genes may result in excessive metabolic burden, which could be achieved within the bandit framework by combining different goals into a multi-objective method [45]. We constrain our design space over 6-core parts of RBS sequences in this study, the design space will increase exponentially when designing a longer sequence. The computational complexity of calculating and sorting acquisition functions (e.g. UCB scores) can be reduced by discretizing the space adaptively and hierarchically [46, 47].

In the Learn phase, our approach has correctly identified the correlation of high guanine content in the RBS with high TIR. We have achieved this despite the relatively low Spearman scores for our predictions. This observation of useful recommendations despite low prediction scores corroborates recent evidence from other studies [20, 48]. Finally, our predictor Gaussian Process regression model, compared with previously described calculators using a deterministic thermodynamic approach, is able to show the uncertainty of the predictions, which can be used by our bandit algorithm to give better recommendations.

In this study we have limited the number of design rounds to four. There were a number of reasons for this, including limitations on time and money, but also the results obtained showed that we have achieved our goal of generating very strong RBS designs. There is a possibility that increasing the number of experimental rounds would enable us to improve the results further, however this has to be put in the context of limited resources. For example, scanning the whole space would surely achieve the best results, i. e. would enable us to find the strongest possible RBS, but that would require unreasonable use of resources. Compared to solutions like the one reported by Hollerer [27], our solution can be used when a high-volume method for data-generation is not available, while still providing the required results (optimised part).

There are still open questions that need to be addressed for applying machine learning in synthetic biology. Firstly, we would like to understand how we can extract more biologically-important information from the decisions made by our algorithms. We have shown that the algorithms are able to exploit them, but it will be important to create tools that will enable their reliable extraction from the results obtained. Secondly, given the small number of RBS sequences tested, how can machine learning algorithms provide more accurate predictions and uncertainty measurements? Thirdly, the generalisability of the method is unknown. We believe that the method described here would be useful for designing other small genetic parts, but the complexity of the task quickly increases with the size of the analysed sequence, so the method’s applicability might be impacted at some point. Similarly, the re-usability of the obtained data set is currently unknown. The TIR of an RBS is dependent on its genetic context, but our Hamming distance and TIR impact of nucleotides outside of the core (Figures 4B and S2 respectively) studies indicate that as long as the changes in the genetic context are small the obtained data set could serve as a basis for similar design efforts. For example, the data could be used to teach the algorithm for the first round of new designs, which in turn could decrease the number of rounds required to obtain the required characteristics. Finally, the practical optimal exploration-exploitation balance between rounds and samples is still an open question.

We have found our approach of bringing machine learning and synthetic biology experts together very fruitful. Synthetic biology is promoting standardised and normalised testing in biology and naturally pairs with machine learning, which can leverage the high quality biological data sets generated when the correct design rules are observed. The addition of machine learning to synthetic biology also adds an additional layer of scrutiny to the generated data sets through the advanced statistical methods that can be used to design and analyse the experiments. On top of that, the use of automation has helped us to produce more reliable results, which gave us the required confidence in our predictions and recommendations. We envision that pairing machine learning with high-throughput automation will keep delivering a high number of good quality data sets and improved methods for biological engineering.

In future we hope to extend the algorithm to other more complicated genetic elements, including promoters and terminators. However, it is important to reiterate that the experimental space grows exponentially with the number of examined positions, so the space becomes increasingly hard to cover with experiments. To solve this problem, different algorithms or experimental techniques might be needed, but the general workflow can be reused.

## 4 Materials and Methods

### 4.1 Laboratory experimental design

#### 4.1.1 BUILD: Construction of genetic devices

##### Plasmid Design

The pBbB6c-GFP plasmid has been used for all our designs. This plasmid contains the GFP mut3b CDS, expression of which can be induced by the addition of isopropyl *β*-D-1-thiogalactopyranoside (IPTG) (Merck, Darmstadt, Germany, catalogue no. I5502). The original RBS for the GFP CDS was replaced using a combination of PCR and isothermal assembly. Primer sequences and the assembly strategy were generated using the Teselagen DESIGN software (Teselagen Biotechnology, San Francisco, CA).

##### PCR

PCR amplification of the cloning inserts was done using Q5 High-Fidelity 2X Master Mix (NEB, Ipswich, MA, catalogue no. M0492L). 20 *μ*L reactions were prepared by dispensing 1 *μ*L of each 10 *μ*M reverse primer into the wells of a 96-well PCR plate using the Echo liquid handler (Beckman Coulter, Brea, CA). A mastermix consisting of polymerase premix, plasmid DNA template (pBbB6c, 5-10 ng per reaction), and the single 10 *μ*M forward primer was prepared and dispensed using the FeliX liquid handler (Analytik Jena, Jena, Germany) or electronic multi-channel pipette. Reactions were run using Touchdown PCR or standard PCR cycling methods in C1000 thermal cyclers (Bio-Rad, Hercules, CA). Capillary electrophoresis of PCR products was performed using the ZAG DNA Analyzer system (Agilent Technologies, Santa Clara, CA). 2 *μ*L of each PCR reaction was electrophoresed using the ZAG 130 dsDNA Kit (75-20000 bp) or ZAG 110 dsDNA Kit (35-5000 bp) (Agilent Technologies, catalogue no. ZAG-110-5000; ZAG-130-5000). ZAG sample plates were prepared using the Sciclone G3 liquid handler (Perkin Elmer, Waltham, MA). ProSize Data Analysis Software (Agilent Technologies) was used to generate gel images from the sample chromatograms, and amplicon sizes were estimated by reference to the upper and lower DNA markers spiked into each sample and a DNA ladder run in well H12 of each sample plate.

##### Isothermal DNA Assembly

Constructs were assembled using NEBuilder HiFi DNA Assembly Master Mix (NEB, catalogue no. E2621L). Reactions consisting of approximately equal amounts of the common fragment and the variable fragment were prepared using the FeliX liquid handler or electronic multi-channel pipette, to a final volume of 5 or 10 *μ*L. Assemblies were run in the thermal cycler for 1 hour at 50°C, followed by an infinite hold step at 4°C. Finally, samples were incubated with of 50 nL of DpnI (NEB, catalogue no. R0176S) at 37°C for 90 minutes to degrade any residual template DNA.

##### *E. coli* transformation

The DH5α cell line (Thermo Fisher Scientific, Waltham, MA, catalogue no. 18265017) was made chemically competent using the Mix & Go *E. coli* Transformation Kit & Buffer Set (Zymo Research, Irvine, CA, catalogue no. T3001). 20 *μ*L of cells was aliquoted into each well of a cold 96-well PCR plate and stored at −80°C for later use. Plates of cells were thawed on a −20°C cold block before 3 *μ*L of the assembly product was added and mixed using the FeliX liquid handler. Cells were incubated on a cold block for 2-5 minutes before being plated in a 96 (12 x 8) grid on Omnitrays containing LB (BD, Franklin Lakes, NJ, catalogue no. 244610) and 15 g/L agar (Merck, catalogue no. A1296) with 34 *μ*g/mL chloramphenicol (Merck, catalogue no. C1919). Plates were incubated overnight at 37°C. Cells were plated using the FeliX liquid handler.

##### Automated colony picking and culturing

A PIXL colony picker (Singer Instruments, Roadwater, United Kingdom) was used to select individual colonies from the transformation plates using the 490-510 nm (cyan) light filter. Each selected colony was used to inoculate 1 mL of selective medium in a 2 mL square well 96 plate. They were then cultured overnight at 37°C with shaking (300 rpm).

##### Glycerol stock preparation

100 *μ*L of sterile 80% (v/v) glycerol (Chem-Supply, Gillman, Australia, catalogue no. GA010) and 100 *μ*L of overnight culture were combined in the wells of a 96 deep (2 mL) round well plate using the FeliX liquid handler or electronic multi-channel pipette They were then sealed with a 96-well silicone sealing mat and transferred to a −80°C freezer.

##### Sequencing

Strains that gave GFP fluorescence intensity readings similar to that of the original RBS were selected for sequence confirmation by capillary electrophoresis sequencing (CES) (Macrogen, Inc., Seoul, South Korea). The strains transformed with each of the selected constructs were grown to saturation in 5 mL LB medium with chloramphenicol selection (34 *μ*g/mL). Plasmids were extracted from the cultures using the QIAprep Spin Miniprep Kit (QIAGEN, Hilden, Germany, catalogue no. 27106) according to the manufacturer’s instructions. Plasmid concentrations were quantified using the Cytation 5 plate reader with the Take3 Micro-Volume Plate (BioTek, Winooski, VT) and all fell in the range of 100-200 ng/μL. Samples of 20 *μ*L of undiluted plasmid DNA were sequenced using a single primer (5’-CGATATAGGCGCCAGCAA-3’) that binds approximately 150 bp upstream of the RBS. Reads were aligned with the template sequence in the Teselagen software.

#### 4.1.2 TEST: Culture analysis

##### Test strain culture

Overnight cultures (six biological (restarted every time from glycerol stock) replicates for each batch) were started by inoculating 1 mL of LB medium supplemented with 34 *μ*g/mL chloramphenicol with 2 *μ*L of the glycerol stock in a 96 deep (2 mL) round well plate, using the FeliX liquid handler or electronic multi-channel pipette. Cultures were incubated at 37°C with shaking (300 rpm) for 17 hours. The following morning, 20 *μ*L of each overnight culture was added to 980 *μ*L of fresh selection medium and these cultures were grown at 37°C with shaking in a 96 deep (2 mL) round well plate. After 90 minutes, cultures were induced with IPTG to a final concentration of 0.5 mM. This was done by transferring 1.0 *μ*L of 0.1 M IPTG to each well of a flat-bottom clear polystyrene 96-well plate using the Echo liquid handler, then adding 300 *μ*L of culture to each well using an electronic multi-channel pipette.

##### Microplate spectrophotometry

The plates were monitored in the Cytation 5 microplate reader immediately after the addition of IPTG. Cytation 5 acquisition and incubation/shaking settings were as follows: length of run: 8 hours; interval: 10 min; continuous orbital shake at 237 rpm and slow orbital speed; excitation wavelength: 490/10 nm; emission wavelength: 515/10 nm; bottom read; gain: 60; read height: 7 mm; read speed: Sweep.

### 4.2 Machine learning experimental design

Two types of machine learning algorithms have to be applied to drive the experimental design workflow shown in Figure 1. One type of machine learning algorithm is a prediction algorithm (**LEARN**), which helps us learn the function of TIR with respect to RBS sequence. The other type of machine learning algorithm is a recommendation algorithm (**DESIGN**), which recommends RBS sequences to query (test) in each round (in the sense of a single pass through the DBTL cycle) based on the predictions from LEARN.

In Round *t*, prediction and design are based on the results obtained in all previous rounds. Our implementation of the machine learning algorithms was tested in Python 3.6 and used the scikit-learn library [49]. In the following paragraphs, we describe our machine learning pipeline (which is applicable to design Rounds 1-3, denoted as Bandit 1-3) in the following order: data pre-processing, prediction and kernels, and finally recommendation. The pipeline for our 0th Round of machine learning designs, denoted as Bandit-0, is reported in Supplementary A.4.

#### 4.2.1 Data pre-processing

For each RBS sequence, we measured the TIR of 6 biological replicates, where TIR is calculated as an averaged sum of GFP fluorescence divided by Optical Density at 600 nm (OD600) of the culture over time (calculated using values from all the measured data points), following the formula:

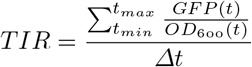

where *t_min_* is the time at which the 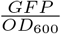 value is at its minimum denoting the start of the log growth phase, *t_max_* is the time at which the logarithmic phase ends (on average, for our samples that was 4 hours from *t_min_* so that was the time point used for TIR calculations for all samples), Δ*t* is the difference between *t_max_* and *t_min_*.

For label pre-processing, we first adjust the TIR values in each round using the round-wise reference values.

The reference value is the TIR of the benchmark sequence that is run in triplicate in each round. Specifically, in Round *t*, we subtract the TIR mean of the benchmark RBS measured in Round *t* from all TIR values measured in the same round, for each replicate separately. We then normalised the data by performing a logarithm transformation and standardisation on the adjusted TIR label for each replicate separately. After normalisation, each replicate has zero mean and unit variance. Furthermore, we also normalised the kernel matrix used for prediction by centring and unit-norm normalisation, which is reviewed in details in Supplementary A.2.1.

#### 4.2.2 Prediction: Gaussian Process Regression with String Kernel

To find RBS sequences with the highest possible TIR score after a total number of rounds *N*, we consider our experimental design problem as a sequential optimisation of an unknown reward function 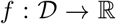, where 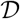 is the set containing all RBS sequence points in the design space, and *f*(**x**) is the TIR score of the 6-base core sequence of the RBS **x** ∈ {*A, C, G, T*}^6^. In each Round t, we choose a set of *m* points 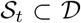 and observe the function value at each point in the selected set 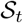, i.e. *y_i_* = *f*(**x**_*i*_) + *ϵ*, for all 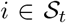, where *ϵ* is the Gaussian random noise with unknown mean and standard deviation.

For the regression model, we have used a Bayesian non-parametric approach called *Gaussian Process Regression (GPR)* [28, 30, 50]. We model *f* as a sample from a *Gaussian Process* 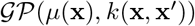, which is specified by the mean function 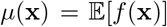 and the kernel function (also called *covariance function*) 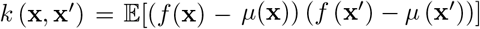. GPR can predict both the posterior mean and standard deviation of the RBS sequences in the design space. The posterior standard deviation represents the level of uncertainty for the prediction.

We review the predictions of GPR in Supplementary A.1.

The choice of kernel function is critical for accurate predictions, since it controls smoothness and amplitude of the function to be modelled. For Bandit designs in Round 0, since we only had access to a limited number of data points from the literature, we chose to use one of the basic string kernels, the *spectrum kernel* [51] to process the core 6 bp and dot product kernel [28] (with one-hot embedding) to process the 7 bp flanking sequences both upstream and downstream of the core sequence. To represent the RBS sequences in subsequent rounds, we use the *weighted degree kernel with shift* (WDS) [52, 53] to specify the kernel function of *GP*. WDS is a type of a string kernel, which takes two sequences as inputs and outputs a scalar value which represents the similarities between the two sequences. WDS kernel does this by counting the matches of substrings of a certain length (i.e. kmers) that constitute the sequence. The maximum substring length is specified by *ℓ*. The WDS takes into account the positional information by counting substrings starting from different positions, where the start position is specified by *l*. Additionally, the WDS kernel considers the shifting of substrings, with the maximum shift specified by *s*. This could be useful when there is a shift between two sequences. For example, two sequences A**CCTGA** and **CCTGA**A are in 1-shift.

We now define WDS kernel. Let 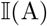 be the indicator function, which equals 1 if *A* is true and 0 otherwise. Then 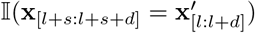 indicates whether two substrings of length *d* match, between **x** starting from position *l* + *s* and **x**′ starting from position *l*. This is similarly done for 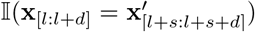. By having these two terms considering substrings of two sequences with starting positions differing by *s* characters, the WDS can measure shifted positional information. When *s* = 0, the kernel function counts the matches with no shift between sequences. Let **x**, **x**′ be two RBS sequences with length *L*, the WDS kernel is defined as

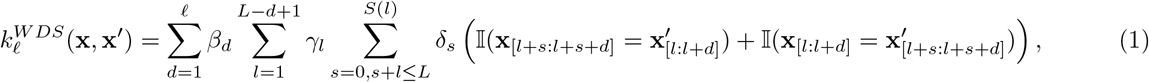

where 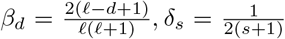, *γ_l_* is a weighting parameter over the position in the sequence, where we chose to use a uniform weighting over the sequences, i.e. *γ_l_* = 1/*L* (*L* = 20 in our design).

*S*(*l*) determines the maximum shift at position *l*. Furthermore, we normalise the kernel with centring and unit-norm in terms of all sequences in the design space. The hyperparameters for kernel, including maximum substring length *ℓ*, maximum shift length *S*(*l*), and the standard deviation of GP noise *α* were chosen based on 10-repeat 5-fold cross validation.

#### 4.2.3 Recommendation: Upper Confidence Bound multi-armed Bandit algorithm

To recommend the RBS sequences to query in the next round, we have used the *Upper Confidence Bound (UCB)* batch algorithm [9], which provides *exploration-exploitation balance*. On one hand, UCB exploits the function in terms of the design space, that is to pinpoint sequences that are believed to have high labels (i.e. high predicted mean); on the other hand, UCB explores the design space where we have little information and sequences have a chance to have high labels (i.e. high predicted standard deviation). More precisely, the UCB algorithm selects RBS sequences 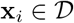 with the maximum upper confidence bound at Round *t*, i.e.

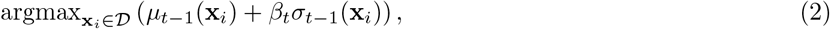

where *β_t_* is a hyperparameter balancing the exploration and exploitation, *μ_t_*(**x**_*i*_), *σ_t_*(**x**_*i*_) are the predicted mean and standard deviation at Round *t* for the sequence **x**_*i*_. We call *μ*_*t*−1_(**x**_*i*_) + *β_t_σ*_*t*−1_(**x**_*i*_) the *UCB score* of sequence **x**_*i*_ at Round *t* − 1.

Since experimentally labelling sequences is time-consuming, it is unrealistic to recommend sequences sequentially (i.e. one-by-one) and then wait for the label to be tested and used to improve the model. Instead, we can recommend RBS sequences in a batch of size *m*. One naive approach is to However, this approach may end up recommending similar sequences in the same local maximum (e.g. *x* = 2, *x* = 2.5 in this example). Since GPR assumes similar sequences would have similar labels (e.g. by knowing *x* = 2 we can gain information about *x* = 2.5 as well), we prefer to not waste time and money on labelling sequences with high similarities in the same batch. To counter this, we use a batch recommendation strategy that is designed to avoid recommending such highly similar sequences in the same batch, described below.

A key property of Gaussian Process regression is that the predicted standard deviation depends only on features, not on the labels. One can make use of this property to design a batch upper confidence bound (BUCB) algorithm [29]. The strategy here is to recommend RBS sequences one-by-one by sequentially adding the newly recommended RBSs’ predicted means (without testing them) into the training data and updating UCB scores.

As illustrated in Figure 5B, the algorithm recommends the data point with maximum UCB score based on the predictions over initial 5 observations. We then add the recommended data point (*x* = 2) into the training data set with the predicted mean of that point as the label (note it is not the true label), and update the predicted standard deviation, then we finally update the UCB scores. The second data point is then recommended based on the new UCB scores. Figure 5C shows that since we assume we have observed *x* = 2, the new predicted standard deviation of the data points in design space around *x* = 2 decreases, so instead of recommending a similar data point *x* = 2.5, the algorithm recommends *x* = 8, which is in another local maximum design area. In this way, the batch recommendation can potentially cover more local maximum areas than the sequential design. In summary, the exploration efficiency is improved since the recommended sequences in one batch will tend to be in different design areas so that the information gain is maximised. In our design, we have set the batch size to 90, to fit the experimental batch. Finally, we set *β_t_* = 2 for Round 0-3 and *β_t_* = 0 for the last round to allow for more exploitation.

**Figure 5:**
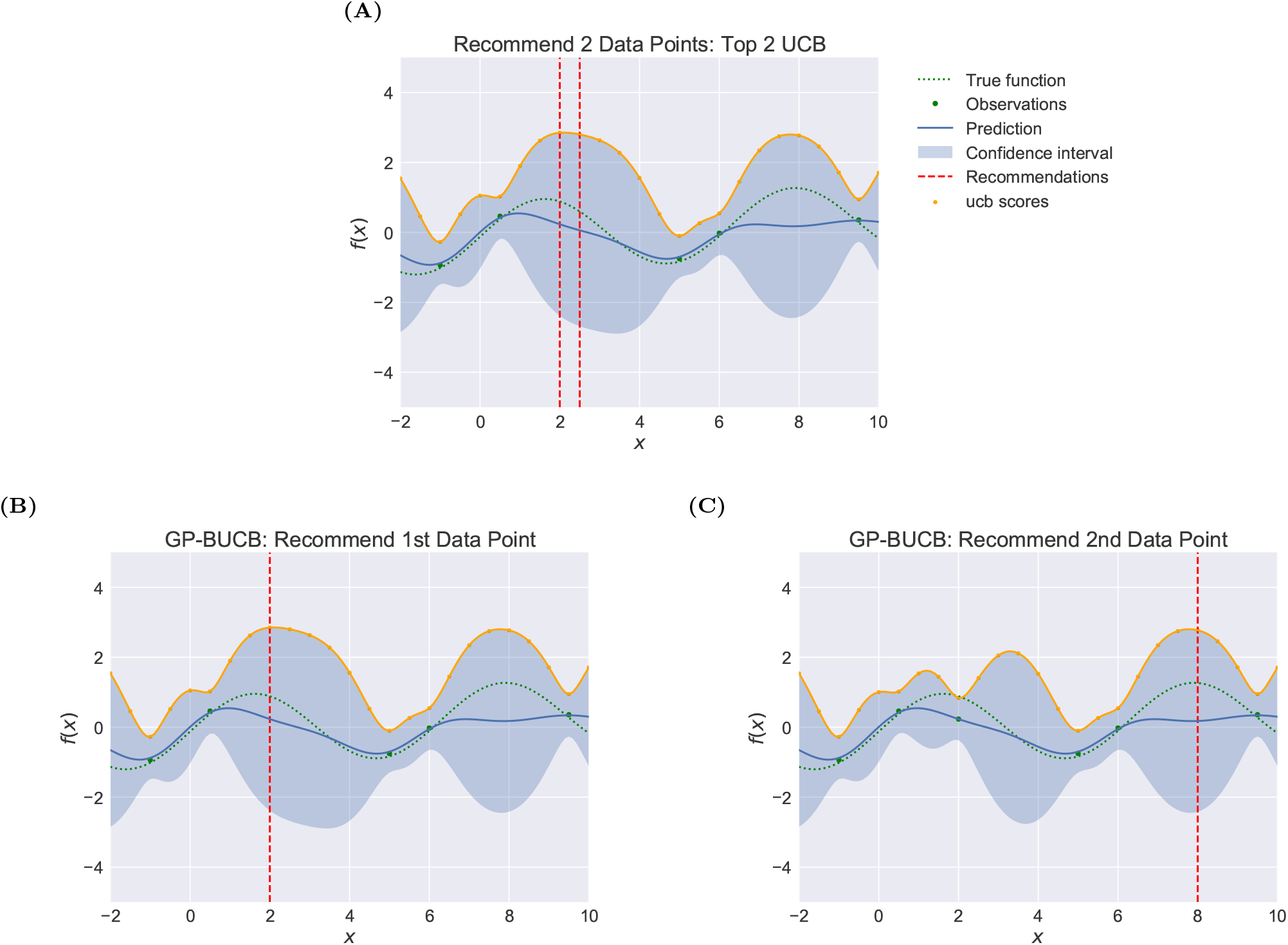
Batch Recommendation Illustration. We use the batch size of 2, with 5 initial observations. The design space is 24 uniformly distributed points in the range [-2,10], i.e. −2, −1.5, −1, …, 9.5, 10. The confidence intervals are shown with predicted mean ±1.96 standard deviation. (A) Top UCB recommendations. The recommendations are 2 data points with top UCB scores, chosen with GP predictions. (B)(C) Batch UCB recommendations. (B) shows the first recommended sequence, (C) shows the new predicted confidence interval and the second recommendation based on that updated interval.

## Code, data and material availability

All code and data required to reproduce the results are available at Github (San Francisco, CA): https://github.com/mholowko/Solaris/tree/master/synbio_rbs. All the processed and raw data are included in the repository. Sequences of plasmids and oligos and assembly reports used in this study are available in the supplementary information as a separate file. The pBbB6c plasmid is available on Addgene (Watertown, MA). Other strains and plasmids are available on request from the authors; MTAs will need to be negotiated between the parties.

## Contributions

Zhang M. and Ong C. S. designed and implemented the machine learning algorithms and workflow. Holowko M. B. and Hayman Zumpe H. have designed and performed the laboratory experiments. Holowko M. B. and Ong C. S. conceived and planned the project. All authors analysed the data, contributed to and reviewed the manuscript.

## Acknowledgments

The authors would like to acknowlege CSIRO’s Machine Learning and Artificial Intelligence, and Synthetic Biology Future Science Platforms for providing funding for this research. The authors would also like to thank CSIRO BioFoundry for help with performing the experiments.

## Supplementary

### A Machine Learning Methods

In this section, we describe the machine learning methods in details.

#### A.1 Gaussian Process Regression

A *Gaussian process* is a collection of random variables, any finite number of which have a joint Gaussian distribution. We define mean function *μ*(**x**) and covariance function *k*(**x**, **x**′) of a real process *f*(**x**) as

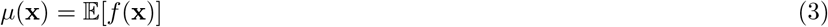

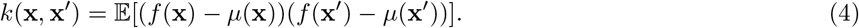

A Gaussian process is specified by its mean function and covariance function as 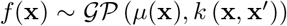. We consider the case where the observations are noisy, i.e. (*X, y*) = {(**x**_*i*_, *y_i_*)|*i* = 1,…, *n*}, where label *y_i_* = *f*(**x**_*i*_) + *ϵ* with 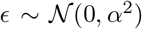. The Gaussian noise is independent identically distributed, and the covariance of the prior on the noisy observations is then cov (*y_p_, y_q_*) = *k*(**x**_*p*_, **x**_*q*_) + *α*^2^*δ_pq_*, where *δ_pq_* is a Kronecker delta which is one if *p* = *q* and zero otherwise. It is equivalent to a diagonal matrix *α*^2^*I* added on the kernel matrix evaluated on the training points.

For *n*_*_ test points *X*_*_, we assume the prior over the functions values as a random Gaussian vector 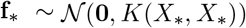. Then the joint distribution of the observed target values and the function values at the test points under the prior as

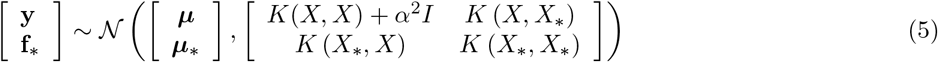

where ***μ***, ***μ***_*_ are mean vectors for **y**, **f**_*_ respectively, *K*(*X, X_*_*) denotes the *n* × *n*_*_ covariance/Kernel matrix evaluated at all pairs of training and testing points, similarly for other kernel matrices. Then the posterior of the test points (i.e. predictive distributions) is given by the conditional distribution 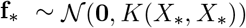, where

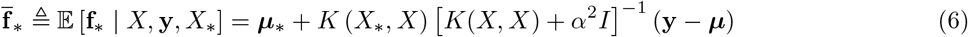

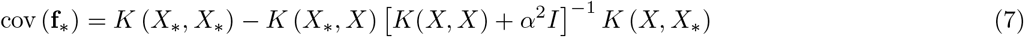

For noisy test targets **y**_*_ = *f*(**x**_*_) + *ϵ* with 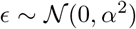, we can compute the predictive distribution by adding *α*^2^*I* to the variance term cov(**f**_*_) in Eq. (7), i.e. 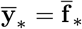, cov (**y**_*_) = cov (**f**_*_) + *α*^2^*I*.

#### A.2 Choices of Kernels

The choice of covariance function is critical for the performance of Gaussian process regression. Except the weighted degree kernel with shift described in Section 4.2.2, we show two closely related string kernels tested in this study below.

- *Spectrum Kernel*.

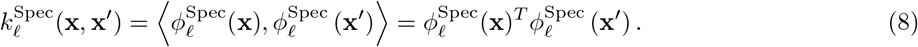

where **x**, **x**′ are two RBS sequences in 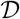 over an alphabet Σ. We denote the number of letters in the alphabet as |Σ|. In our case, Σ = {*A, C, G, T*} and |Σ| = 4. 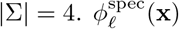 maps the sequence **x** into a |Σ|^*ℓ*^ dimensional feature space, where each dimension is the count of the number of one of the |Σ|^*ℓ*^ possible strings *s* of length *ℓ*. For example, if **x** = *ACACAG* and *ℓ* = 3, then 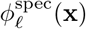 maps **x** into 4^3^ dimensional feature space, where each dimension corresponds to *AAA, AAC, AAT, AAG, ACA*,…, *TTT*. Among those, the dimension corresponds to *ACA, CAC, CAG* are 2,1,1 and others are 0. Let *X*, *X′* be two metrics which include n sequences, and 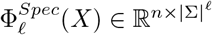, then the spectrum kernel over metrics is

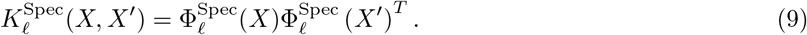
- *Weighted Degree Kernel*, considers positional information. WD kernel counts the match of kmers at corresponding positions in two sequences. For sequences with fixed length *L* and weighted degree kernel considers substrings starting at each position *l* = 1,…, *L*, with 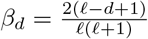,

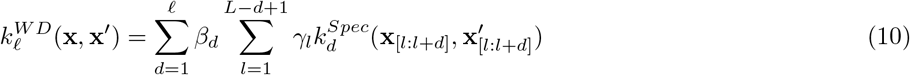

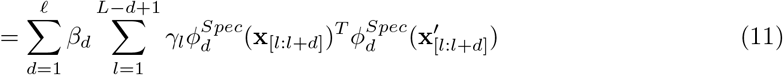

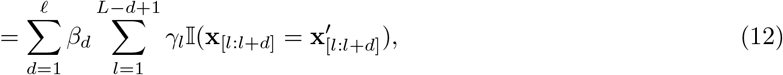

where 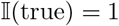 and 0 otherwise.

##### A.2.1 Normalisation of Kernel

As part of data pre-processing, the range of all features should be normalised so that each feature contributes approximately proportionately to the predictive model. The kernel matrix is represented by the inner product of the underlying feature vectors, it needs to be normalised before being used in the downstream regression models. Upscaling (down-scaling) features can be understood as down-scaling (up-scaling) regularizers such that they penalise the features less (more).

Here we consider two approaches for kernel normalisation: centering and unit norm. We will show how to convert the normalisation in terms of feature vectors to normalisation in terms of kernel matrices. As defined before, consider **x**, **x**′ are two RBS sequences in 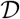 over an alphabet Σ. We denote *ϕ*(**x**) as a column feature vector of sequence **x**, where a feature function 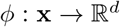, with *d* as the dimension of features. One example of *ϕ* can be 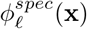 in the spectrum kernel. Recall the corresponding kernel can be defined as *k*(**x**, **x**′) = *ϕ*(**x**)^*T*^*ϕ*(**x**).

Assume there is total of *n* sequences in the data *X*(*n*′ sequences in the data *X*′). We illustrate centering and unit norm normalisation below.

- Centering. Defining the mean vector as 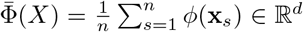, the centered feature vector 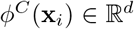 of **x**_*i*_ is

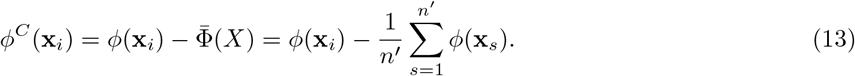 The corresponding centering kernel value between **x**_*i*_ and **x**_*j*_ is then

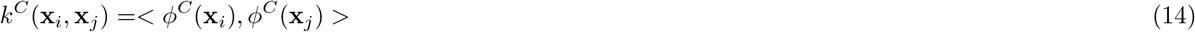

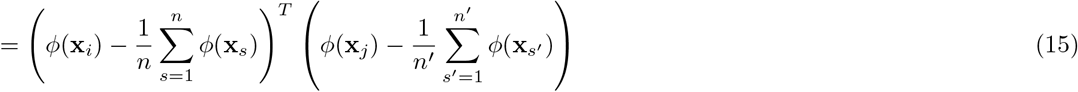

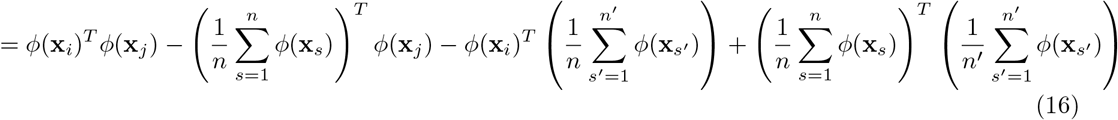

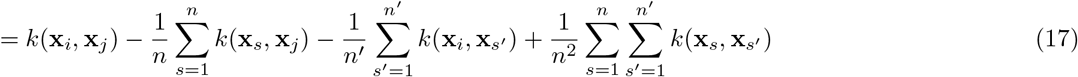
- Unit Norm. Define the (*l*_2_) norm of a feature vector as 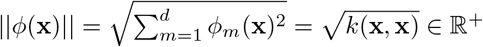, then the unit norm feature vector 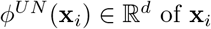 of **x**_*i*_ is

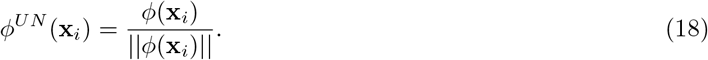 The corresponding unit norm kernel value between **x**_*i*_ and **x**_*j*_ is then

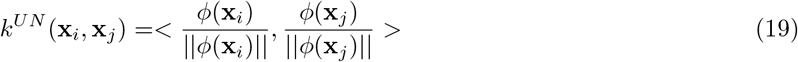

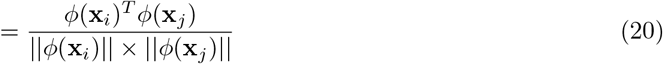

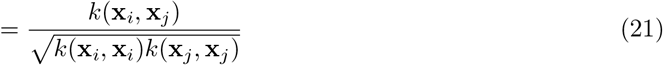
- Unit Variance. After the centering and unit norm normalisation, the kernel matrix is unit variance as well. In the following, we show transformations of the unit variance (with centering) normalisation. Define the variance vector 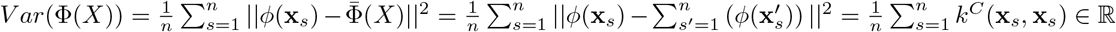, the unit variance feature vector 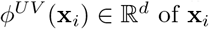 of **x**_*i*_ is

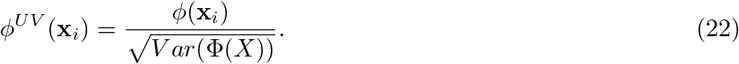 The corresponding kernel representation is

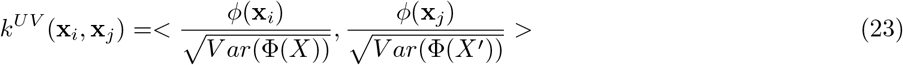

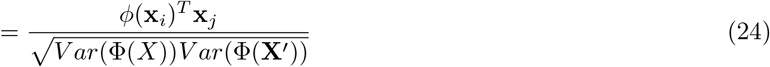

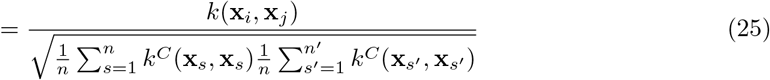 After centering and unit norm, 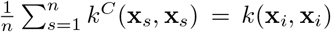, which implies that after centering and unit norm, the kernel matrix is already unit variance normalised.

For the Gaussian Process regression, we make of use of two kernel matrices: the kernel function between the training data itself, i.e. *K*(*X_train_, X_train_*); and the kernel function taking the training data and testing data as inputs, i.e. *K*(*X_test_, X_train_*). We will state two ways of normalisation those two kind of matrices:

- Normalise training and testing data separately. This approach is preferred for most of the machine learning algorithms since it follows the rule that we have no information about testing data while training. Then for centering, one should subtract the mean vector over the training data for both kinds of matrices. For unit norm normalisation, when one calculates *K^UN^*(*X_test_, X_train_*), the two terms inside of square root: *k*(**x**_*i*_, **x**_*i*_) is taken from *K*(*X_test_, X_test_*)[*i, i*], and *k*(**x**_*j*_, **x**_*j*_) is taken from *K*(*X_train_, X_train_*)[*j, j*].
- Normalise training and testing data together, i.e. normalise *K*(*X_train+test_, X_train+test_*), then extra the parts we need from the normalised matrix. This approach is suitable in a case where one already knows the whole of testing features. For centering, one should subtract the mean vector over the whole matrix Φ(*X_train+test_*). The unit norm normalisation is the same as in the previous case.

For our experiment, we fix the design space before training, i.e. the testing features are already known before testing. So we choose to normalise the kernel matrix over the training and testing data together (for Round 2-3, normalise over all RBS sequence in the design space), by first applying centering and then unit norm normalisation.

#### A.3 Batch Recommendation

We consider recommending sequences in batch and using Gaussian Process Batch Upper Confidence Bound (GP-BUCB) algorithm [29]. We show technical details of GP-BUCB in the following. With batches of size *B*, the feedback mapping *fb*[*t*] =| ⌊(*t* − 1)/*B*⌋*B*, i.e.

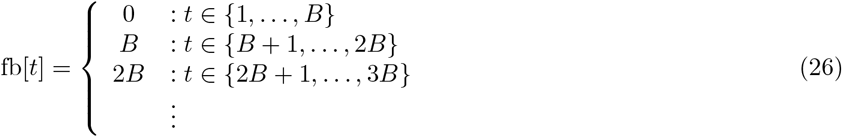

A key property of Gaussian Process regression is that the predictive variance in Eq. (7) only depends on observed points (i.e. features), but not on the labels of these observed points. So one can compute the posterior variance without actually observing the labels. The GP-BUCB policy is to select sequences that

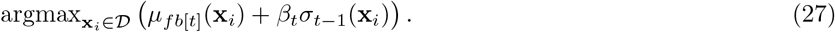

And only update *y_t′_* = *f*(***x**_t′_*) + *ε_t′_* for *t′* ∈ {*fb*[*t*] + 1,…, *fb*[*t* + 1]} at the end of each batch (*fb*[*t*] < *fb*[*t* + 1]). This is equivalent to sequential GP-UCB with *hallucinated observations **y***_fb[*t*]+1:*t*−1_ = [*μ*_fb[*t*]_ (***x***_fb[*t*]+1_),…, *μ*_fb[*t*]_ (***x***_*t*−1_)], while the posterior variance decreases.

#### A.4 Bandit-0 Design

The design of Round-0 is based on the literature data [36]. We first normalise the raw TIR to values between 0 and 1. We applied the Gaussian Process Regression with noise parameter *α* = 1*e* − 10. We chose to use one of the basic string kernels, the *spectrum kernel* [51] to process the core 6bp and dot product kernel [28] (with one-hot embedding) to process the 7bp flanking sequences both upstream and downstream of the core sequence. The design size is 60 with UCB parameter *β* = 1.

#### A.5 Addition Results

As stated in Section 2.3, we merely aimed to provide a valid ranking in Learn part due to the small amount of noisy data, which can be captured by the Spearman correlation coefficient. We show *R*^2^ score for the reader’s interest, which quantifies value-based prediction evaluation. The *R*^2^ increases along with rounds increase.

**Table 2:** *R*^2^ score for predictions in Figure 3.

|  | **Train 0** | **Train 0-1** | **Train 0-2** | **Train 80%** |
| --- | --- | --- | --- | --- |
|  | **Test 1** | **Test 2** | **Test 3** | **Test 20%** |
| $R^2$ | 0.067 | 0.226 | 0.213 | 0.269 |
| **Spearman** | 0.269 | 0.468 | 0.546 | 0.479 |

#### A.6 Exploration-exploitation visualisation

In this section, we show our method of exploration-exploitation visualisation displayed in Figure 2C. The UCB score of a sequence **x**_*i*_ at Round *t* is defined as

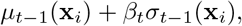

where *β_t_* is a hyperparameter balancing the exploration and exploitation, *μ*_*t*−1_(**x**_*i*_), *σ*_*t*−1_(**x**_*i*_) are the predicted TIR and standard deviation (STD) at Round *t* − 1 for the sequence **x**_*i*_. The sequences with high predicted TIR address exploitation, while the sequences with high predicted STD address exploration. We show the predicted TIR (x-axis) v.s. predicted STD (y-axis) in Figure S1 as orange points.

To compare the predicted TIR and STD in the same dimension (basis), we project the (predicted TIR, predicted STD) pair to the diagonal line (the grey dashline) set between (0,1) and (1,0) coordinates. The slope of the diagonal line is selected to weight the predicted TIR and STD equally. Each point on the diagonal then, corresponds to an orthogonal projection (known as *vector projection*) of an orange point. As a result, the higher the predicted TIR the closer the point is to the (1,0) coordinate, and the higher the predicted STD the closer the point is to the (0,1) coordinate. Finally, we colour those projected points using the RdBu spectrum, where the points with high STD (exploration) are shown in red and those with high mean TIR (exploitation) are shown in blue. Finally, these colours correspond to ones shown in Figure 2C.

**Figure S1:**
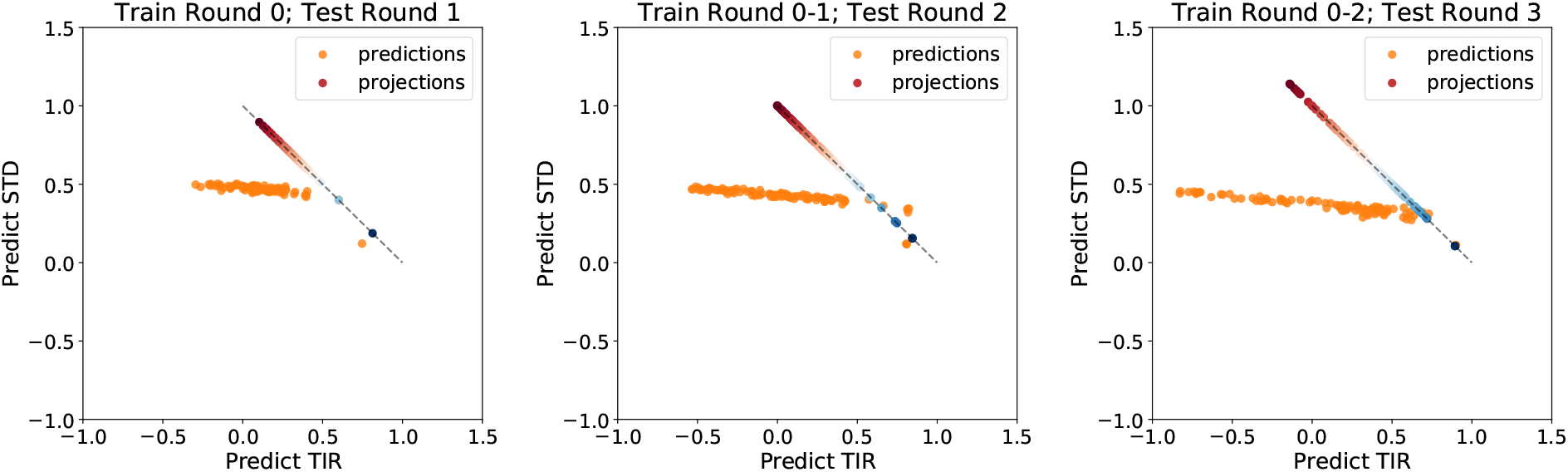
Exploration-Exploitation scores illustration. We show the predictions for each sequence from Round 1-3 in orange (with the same settings as Figure 3) and their corresponding projections to (0,1), (0,1) diagonal coloured in RdBu spectrum. We explain in details how the colours are generated in Section A.6.

### B Supplementary Plots

**Figure S2:**
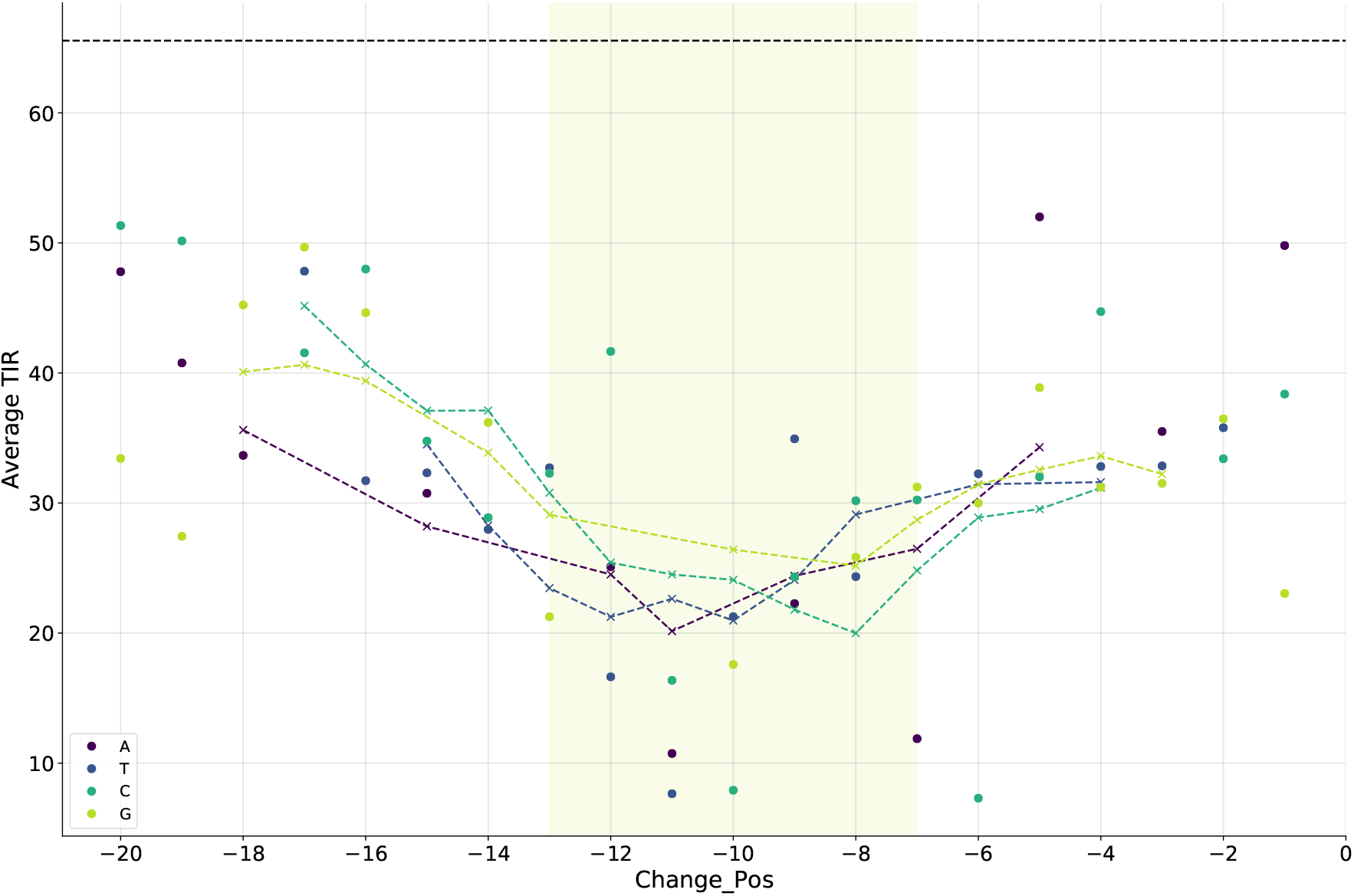
Comparison of base change impact on TIR in core versus non-core region. The core region is highlighted in light green and the lines are rolling averages (in this case mean of the 5 points centred over the indicated position) for each base. The top dotted line shows the TIR for the benchmark sequence, where dots represent a change at a given position to a given base, which is colour coded. Since only changes from the original base at each position are shown, the dotted lines start and end at different positions depending on how many changes to the respective base are at positions adjacent to the indicated one. The value of Welch’s t-test between the mean TIR in core and non-core groups is −4.8780 with p-value < 0.0001 and 34 degrees of Freedom.

**Figure S3:**
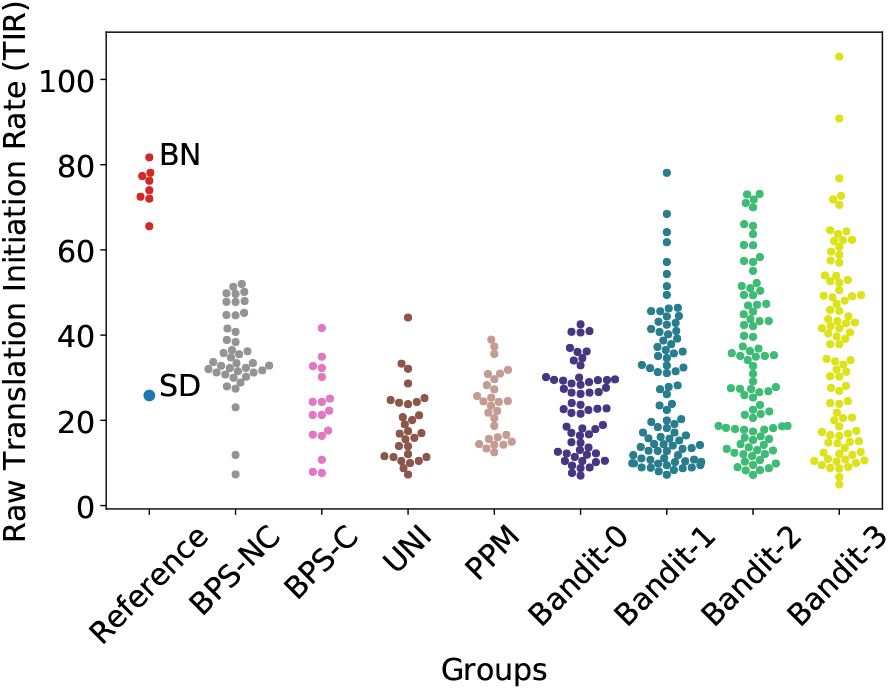
Raw TIRs of RBS groups examined in this study. Raw TIR is calculated as a derivative of GFP fluoresence divided by OD600 of culture over 4h counting from the start of log phase of growth. BPS-NC: base-by-base changes in the non-core region. BPS-C: base-by-base changes in the core region. UNI: Randomly generated sequences with uniform distribution. PPM: Randomly generated sequences with distribution following the PPM for all natural RBS in *E. coli*. Bandit-0/1/2/3 - Bandit algorithm generated results for Round 0, 1, 2 and 3 respectively. SD - Shine-Dalgarno sequence. BN - benchmark sequences for all plates.

**Figure S4:**
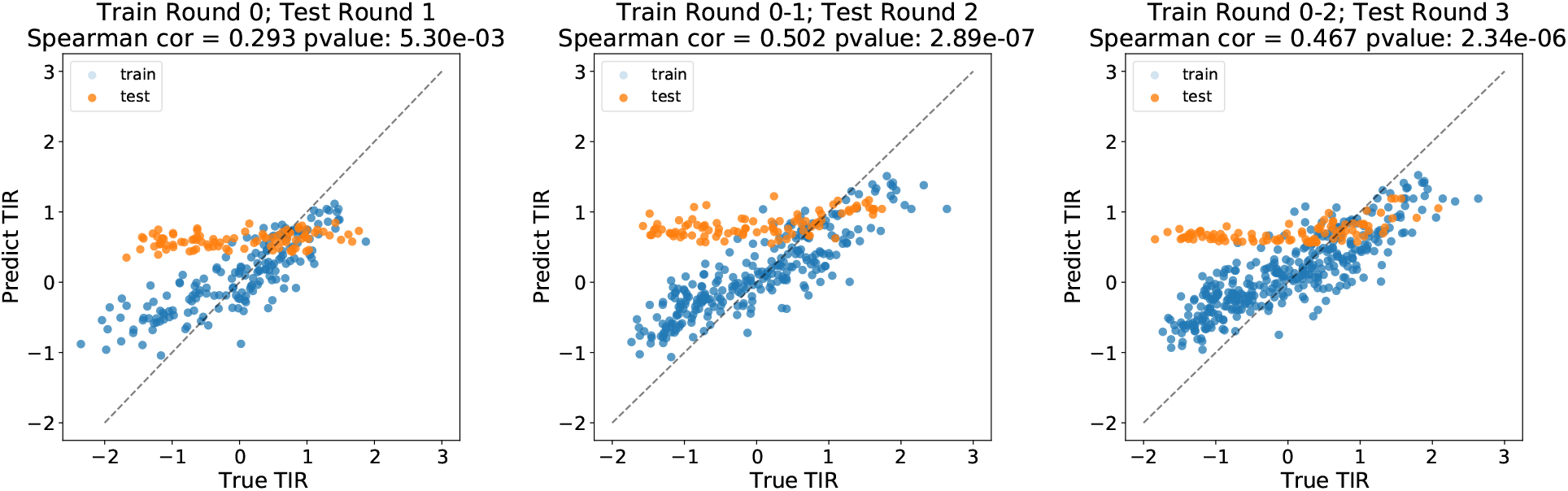
Predictions in design pipeline in Round 1-3. For Round 1, the kernel matrix is normalised based on known RBS sequences in Round 0. For Round 2-3, the kernel matrix is normalised based on all RBS sequences in the design space. Compared to the results shown in Figure 3, here we have applied kernel normalisation described in Section A.2.1. Although the precise value prediction looks different from Figure 3, the rankings provided in two figures are similar, which can be reflected by the similar Spearman correlation coefficients.

**Figure S5:**
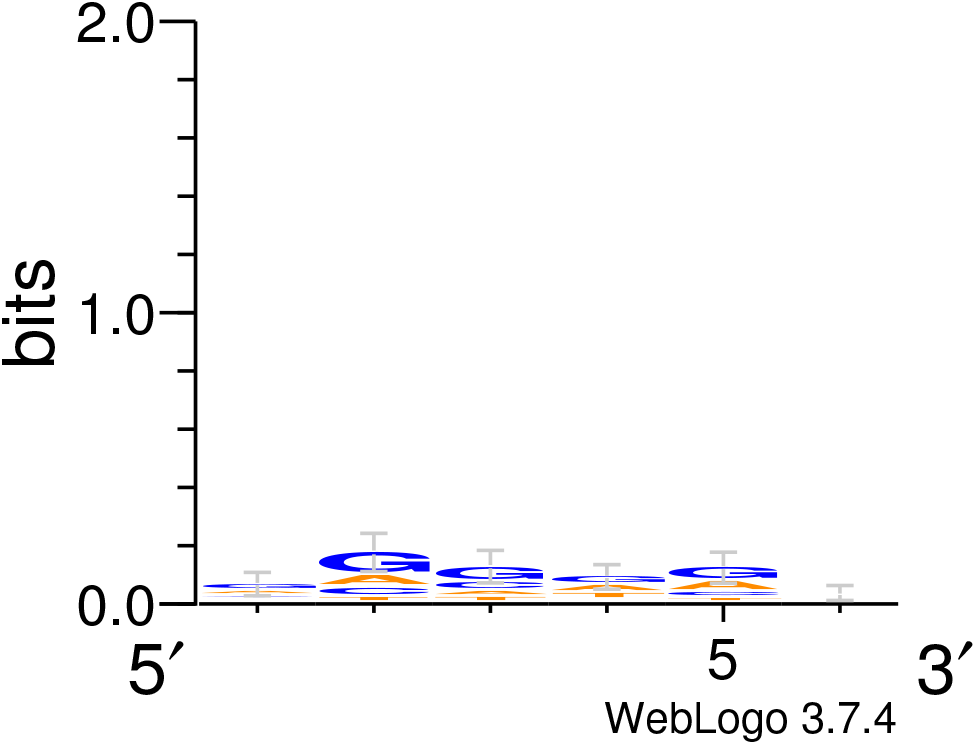
Sequence logo for all tested sequences. Compared to Figure 4A there are no significant biases at each position.

**Figure S6:**
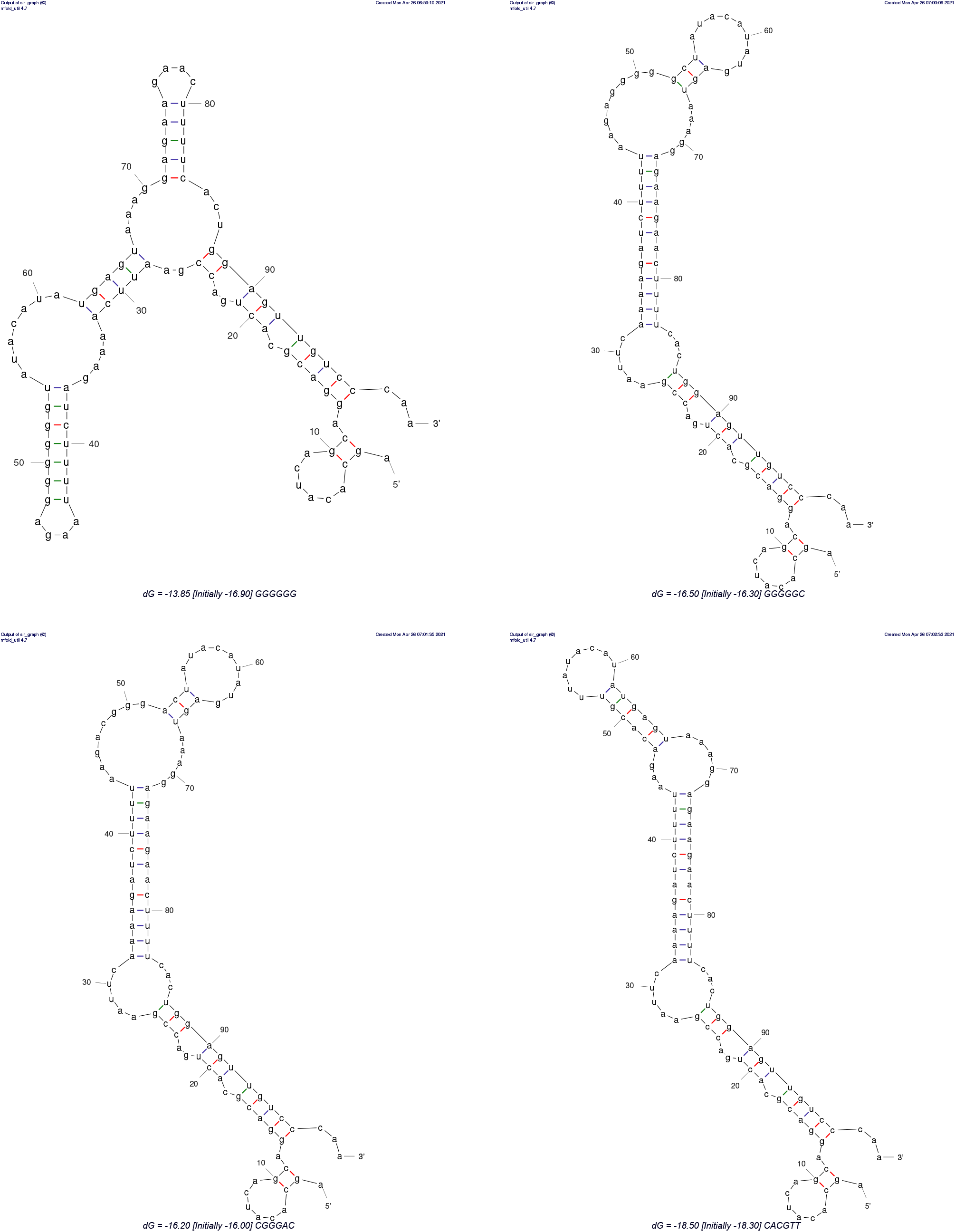
Folding predictionds for four different RBS. Here we show the predicted, energetically most favourable structures for four of our RBS with the following cores: GGGGGG, GGGGGC, CGGGAC, CACGTT and their immediate upstream and downstream background sequence. There is no discerbile and consistent difference between strong ones (the first two) and weak ones (the last two).

**Figure S7:**
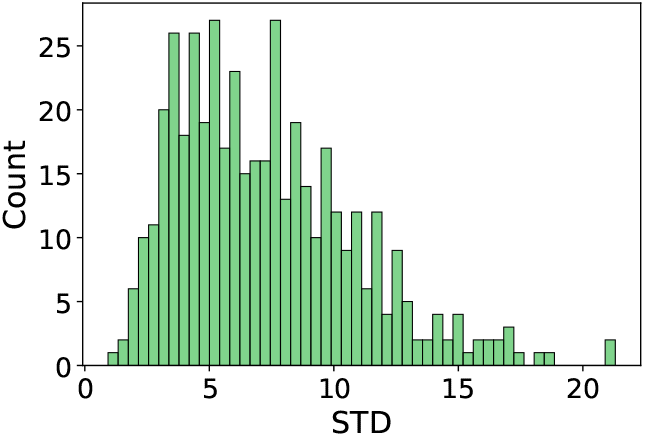
Histogram of standard deviations. For each RBS sequence, we tested 6 biological replicates. This plot shows the histogram of the STD of raw TIR values of the 6 replicates for each RBS sequence.

**Figure S8:**
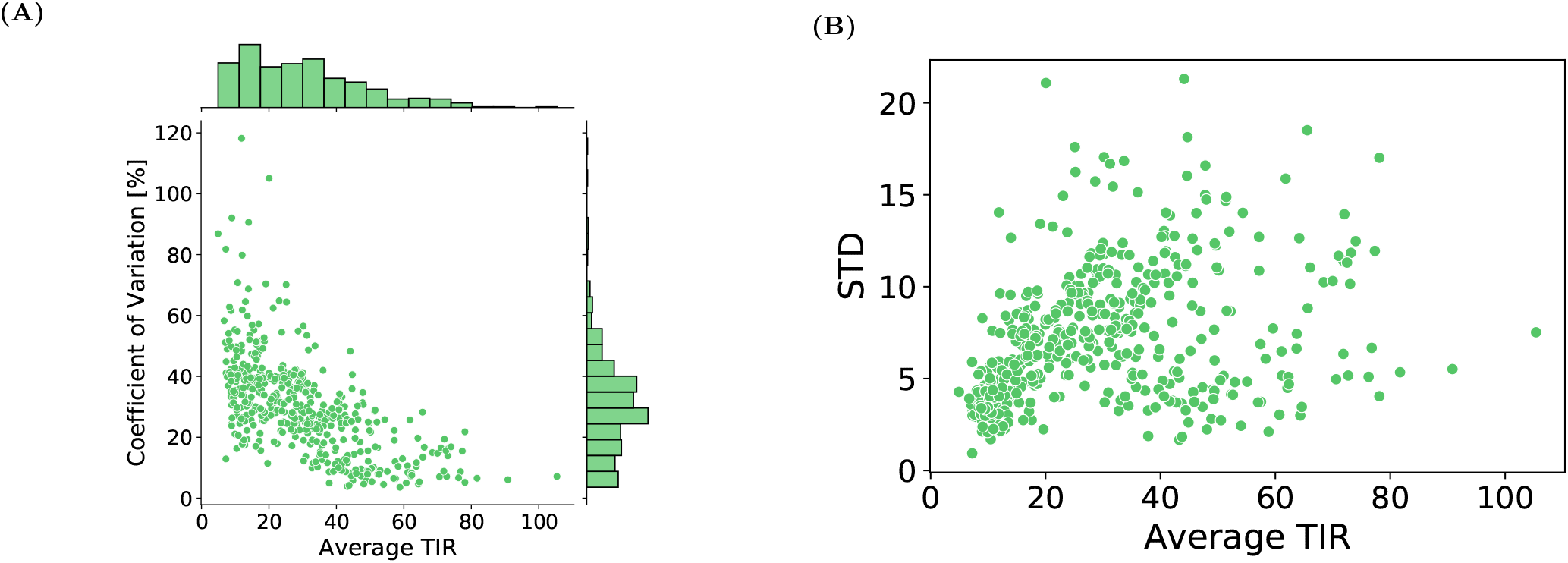
Distribution of standard deviation of tested samples. A) Joint plot showing the distribution of relative error calculated as the average TIR from 6 replicates divided by the given samples standard deviation (coefficient of variation). B) Scatter plot showing the standard deviation for each RBS plotted against its averaged TIR. Standard deviation and average TIR is given in terms of raw number.

